# A homodimeric aptamer variant generated from ligand-guided selection activates T-cell receptor cluster of differentiation three complex

**DOI:** 10.1101/2020.05.18.102145

**Authors:** Lina Freage, Deana Jamal, Nicole Williams, Prabodhika R. Mallikaratchy

**Author notes:** To whom correspondence should be addressed: Prabodhika Mallikaratchy, Department of Chemistry, Lehman College, The City University of New York, 250 Bedford Park West, Bronx New York, NY 10468.

## Abstract

Recently, immunotherapeutic modalities with engineered cells and monoclonal antibodies have been effective in treating several malignancies. However, growing evidence suggests that immune-related adverse events (irAE) lead to severe and long-term side effects. Most iRAEs involve prolonged circulation of antibodies. To address this problem, nucleic acid aptamers can serve as alternative molecules to design immunotherapeutics with high functional diversity and predictable circulation times. Here, we report the first synthetic prototype consisting of DNA aptamers, which can activate T-cell receptor cluster of differentiation 3 (TCR-CD3) complex in cultured T-cells. We show that activation potential is similar to that of a monoclonal antibody (mAb) against TCR-CD3, suggesting the potential of aptamers in developing efficacious synthetic immunomodulators. The synthetic prototype of anti-TCR-CD3ε, as described herein, was designed using aptamer ZUCH-1 against TCR-CD3ε, generated by Ligand Guided Selection (LIGS). Aptamer ZUCH-1 was truncated and modified with nuclease-resistant RNA analogs to enhance stability. Several dimeric analogs with truncated and modified variants were designed with variable linker lengths to investigate the activation potential of each construct. Among them, dimeric aptamer with approximate dimensions similar to those of an antibody showed the highest T-cell-activation, suggesting the importance of optimizing linker lengths in engineering functional aptamers. The observed activation potential of dimeric aptamers shows the vast potential of aptamers in designing synthetically versatile immunomodulators with tunable pharmacokinetic properties, expanding immunotherapeutic designs with the use of nucleic acid-based ligands such as aptamers.

## Introduction

Recently, mAbs have been successfully utilized in designing immunotherapeutic modalities(1,2). These modalities are aimed at treating various types of malignancies. However, an increasing amount of evidence suggests that mAb-based treatments against immunological markers lead to various autoimmune diseases and liver-specific toxicity in humans(3,4). These side effects arise from prolonged circulation times of mAbs, owing to their relatively high molecular weights, and this characteristic has been a challenge in all types of mAb-based therapeutic strategies(5). To address this problem, nucleic acid aptamers, considered as synthetic mAbs, could offer alternative molecules to design immunotherapies(6,7). In recent years, the potential of aptamers as immunostimulatory agents has been explored. For example, dimeric aptamers have been developed against 4-1BB expressed on surface of activated mouse T cells and OX40, a stimulatory molecule belonging to the tumor necrosis factor (TNF) superfamily of receptors. Both dimeric aptamers exerted superior stimulatory activity compared to corresponding mAbs against the same targets(8,9). In another study, the immunomodulatory ability of anti-CD28 aptamers has been shown to exert antagonistic or agonistic ability *via* dimerization(10).

While aptamers show tremendous potential in designing various scaffolds for immunomodulation, the challenges lie in the identification of versatile aptamers against key receptors expressed on the immune cell surface. One such critical receptor, which plays a significant role in immune cell activation, is the T-cell receptor cluster of differentiation 3 (TCR-CD3) receptor expressed on human T-cells. TCR-CD3 complex consists of multiple domains(11). Identifying versatile aptamers against such complexes is difficult to achieve through the conventional Systematic Evolution of Ligand by Exponential enrichment (SELEX) process(12,13). Recently, however, we successfully addressed this challenge by introducing a variant of SELEX called Ligand Guided Selection (LIGS)(14-17). The LIGS method is designed to identify aptamers against complex receptor proteins in their native conformation(14-17). Using LIGS, we reported the identification of a panel of aptamers that can specifically recognize the TCR-CD3 complex expressed in human cultured cells and T-cells obtained from healthy individuals(16). So far, the only known ligands against CD3-TCR are mAbs, and to the best of our knowledge, no alternative synthetic ligands are available.

In the last decade, increased interest has shown in truncating aptamers to variants that show enhanced activity(18,19). In fact, we and others have shown that the truncation and dimerization of aptamers resulted in the enhanced binding properties of molecules against cellular targets(18,20,21). However, most dimeric aptamers are confined to enhancing affinity and stability to develop targeting agents to deliver RNA therapeutics. Given the vast potential of aptamers as synthetic immunomodulators, we, herein, explored one such aptamer, termed ZUCH-1, which was identified *via* LIGS as a T-cell activator. We followed post-SELEX modification strategies to systematically truncate the aptamer and modified with nuclease-resistant RNA analogs to enhance structural and nuclease stability. Finally, dimeric analogs were designed with variable lengths to introduce an agent that could activate TCR-CD3 for the design of aptamer-based immunotherapies. Interestingly, we found that linker length between the aptamers plays an essential role in activating TCR-CD3, opening novel avenues in designing functional dimeric aptamers against cell surface receptors.

## Method and Materials

### Cell Culture and Reagents

Jurkat, Clone E6.1 (T lymphocyte), cells were purchased from the American Type Culture Collection (ATCC). Jurkat double-knockout cells generated via CRISPR-Cas9 targeting CD3E and TRAC genes were purchased from Synthego (Redwood City, CA, USA). All cell lines were cultured in HyClone RPMI-1640 (+25mM HEPES, +L-Glutamine) medium supplemented with 100 units/mL penicillin streptomycin 1% (Corning), 1% MEM Non-Essential Amino Acids (Gibco) and 10% fetal bovine serum (Heat Inactivated, Gibco). All cell lines were routinely evaluated on a Flow Cytometer (Becton Dickinson, FACScan) for the expression of CD marker using anti-hCD3ε (PE Conjugated Mouse IgG1, R&D Systems) antibody to authenticate the cell line.

### DNA Synthesis

All DNA reagents needed to perform DNA synthesis were purchased from Glen Research or ChemGenes. All aptamers were chemically synthesized by attaching a 5*’* 6-FAM (Fluorescein) using standard solid-phase phosphoramidite chemistry on an ABI394 DNA synthesizer (Biolytics) using a 1µmol scale. The monomer OSJ-T3-LNA was synthesized using locked analog phosphoramidite bases (Glen Research) to modify the first and the second bases (5’ end) of the aptamer with Locked Nucleic Acid (LNA) while OSJ-T3-OMe was synthesized by using the 2’-OMe-Ac-C-CE and 2’-OMe-U-CE phosphoramidites (Glen Research) to modify the 40^th^ and 41^st^ bases (Glen Research), respectively. OSJ-T3-LNA-OMe was synthesized using both the locked analog phosphoramidite to modify the first and second bases as well as OMe-Ac-C-CE and 2’-OMe-U-CE phosphoramidites to modify the 40^th^ and 41^st^ bases.

Two monomeric OSJ-T3-LNA-OMe molecules were tethered using the spacer Phosphoramidite 18 (Glen Research) to form dimers. Two, four, six, and eight spacer repeats were used to design OSJ-D-2S, OSJ-D-4S, OSJ-D-6S, and OSJ-D-8S dimers, respectively. Synthesized DNA sequences were de-protected according to the base modification employed and purified using HPLC (Waters) equipped with a C-18 reversed-phase column (Phenomenex) and UV detector using 0.1M TEAA as the mobile phase. The random ssDNA molecule and the dimeric random controls were purchased from IDT DNA Technologies.

### Preparation of Solutions

After the purification of all aptamers, the DNA stock solution concentrations were determined by using a UV-Vis spectrophotometer (Thermo Scientific). Sub-stock solutions of 10 µM were prepared for all aptamer molecules by dilution of each of the respective stock solution with RPMI-1640 (+25mM HEPES, +L-Glutamine) medium. The 10 µM solutions were then further diluted in RPMI-1640 (+25mM HEPES, +L-Glutamine) medium to prepare various working solutions. The random control molecules were prepared as described previously.

### Cell Suspension Buffer (CSB)/ Binding Buffer

All binding assays were performed in Cell Suspension Buffer (CSB) composed of HyClone RPMI-1640 (+25mM HEPES, +L-Glutamine) medium containing 200 mg/L tRNA (Sigma-Aldrich), 2 g/L Bovine Serum Albumin (BSA, Fischer Scientific) and 200 mg/L Salmon Sperm DNA solution (Invitrogen).

### Aptamer Folding Conditions

Prior to aptamer binding with the cells, the truncated aptamers prepared in HyClone RPMI-1640 medium were denatured for 5 mins at 95°C and allowed to fold into its secondary structure for 45 mins at 37°C. Dual-modified and dimeric aptamers prepared in HyClone RPMI-1640 (+25mM HEPES, +L-Glutamine) medium were instead denatured for 10 mins at 95°C followed by 15 mins fold interruption on ice then 30 mins at 37°C.

### Cell-binding Assays

#### Affinity assay

Jurkat E6.1 cells were prepared by washing three times with 3 mL HyClone RPMI-1640 (+25mM HEPES, +L-Glutamine) medium prior to aptamer binding to the cells. All aptamers and the random sequences were prepared at an initial concentration of 250 nM from 10 µM sub-stock solutions. Truncated and dual-modified aptamers were denatured and allowed to fold into their secondary structures as previously described. The truncated monomers were initially prepared at a concentration of 250 nM and serially diluted to achieve the desired final concentrations of 62.5 nM, 25 nM, 12.5 nM, 5 nM, 1 nM and 0.5 nM in 150 µL (75 µL of aptamer in RPMI-1640 and 75 µL of 1.5 × 10^5^ Jurkat E6.1 cells in CSB). Dual-modified and OSJ-D-2S, OSJ-D-4S, OSJ-D-6S dimers were initially prepared at a concentration of 25 nM and serially diluted to achieve the desired final concentrations of 12.5 nM, 6.25 nM, 3.13 nM, 1.6 nM, 0.78 nM and 0.02 nM in 150 µL (75 µL of aptamer in RPMI-1640 and 75 µL of 1.5 × 10^5^ Jurkat E6.1 cells in CSB). The OSJ-D-8S dimer was initially prepared at a concentration of 50 nM and serially diluted to achieve the desired final concentrations of 25 nM, 12.5 nM, 6.25 nM, 3.13 nM, 1.6 nM, 0.39 nM and 0.07 nM in 150 µL (75 µL of aptamer in RPMI-1640 and 75 µL of 1.5 × 10^5^ Jurkat E6.1 cells in CSB). Following this, aptamers and cells were incubated at 37°C for 1 hour. After incubation, cells were washed two times with 2 mL RPMI-1640 medium and reconstituted in 250 µL RPMI-1640 medium. Binding of each aptamer was analyzed using flow cytometry by counting 5000 events for a given concentration. As a positive control, Jurkat E6.1 cell lines were incubated with 5 µL of 25 µg/mL anti-hCD3ε antibody (PE Conjugated Mouse IgG_1,_ R&D System) or 2 µL of 200 µg/mL isotype control (PE Mouse IgG_1_, κ, BD Biosciences) for 30 mins on ice, followed by a one-time wash with 2 mL of RPMI-1640 medium and reconstituted in 250 µL RPMI-1640 medium. Binding events were monitored in FL1 Green (515-545 nm) for the aptamer and in FL2 Yellow (565-605 nm; 564-606 nm) for the antibody by counting 5000 events using flow cytometry.

The final amount (moles) of aptamer used to plot the truncated aptamer affinity ranged from 0.075 pmole to 9.375 pmole and 0.003 pmole to 3.75 pmole for the dual-modified, OSJ-T3-LNA-OMe. Dimeric aptamers final amount (moles) ranged from 0.0117 pmole to 3.75 pmole. All experiments were done in triplicates and the specific binding for each concentration was calculated using the equation:

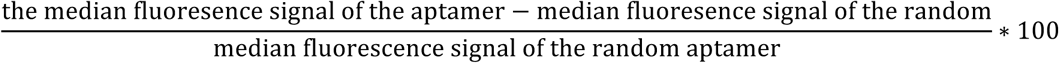

These specific binding values were then used to generate an affinity curve using GraphPad Prism. The Kd values were calculated based on a fit using GraphPad Prism software (one site-specific binding) using the following equation: Y = B_max_*X / (Kd +X), where X is the X-axis, Y is the Y-axis and B_max_ is the maximum number of binding sites.

#### Specificity Assay

Cells were prepared by washing three times with HyClone RPMI-1640 (+25mM HEPES, +L-Glutamine) medium prior to aptamer binding to the cells. Truncated and dual-modified aptamers were denatured and allowed to fold into their secondary structures as previously described. The specific binding was investigated by incubating 75 µL of 0.0375 nmole of each aptamer or random in RPMI-1640 medium with 75 µL of 1.5 × 10^5^ Jurkat E6.1 or TCR-CD3ε double-knockout Jurkat in cells suspension buffer (CSB) resulting in a concentration of 250 nM. Following incubation, the cells were then washed two times with 2 mL of RPMI-1640 medium and reconstituted in 250 µL RPMI-1640 medium. Binding events were monitored in FL1 Green (515-545 nm) for the aptamer using flow cytometry by counting 5000 events for each concentration. As a positive control, Jurkat E6.1 cells were incubated with 5 µL of 25 µg/mL anti-hCD3ε (PE Conjugated Mouse IgG_1,_ R&D System) antibody or 2 µL of 200 µg/mL isotype control (PE Mouse IgG_1_, κ, BD Biosciences) for 30 mins on ice. Cells were then washed one time with 2 mL of RPMI-1640 medium and reconstituted in 250 µL RPMI-1640 medium. Binding events were monitored in FL2 Yellow (565-605 nm; 564-606 nm) for the antibody by counting 5000 events using flow cytometry.

All affinity experiments were done in triplicates, and the specific binding percentages were calculated using the following *equation*:

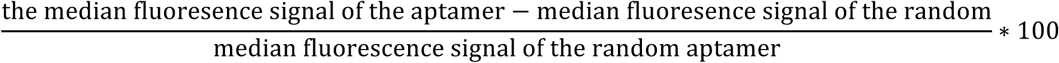

#### Stability of Aptamers

To analyze the stability and nuclease resistance of the dimers, 20 µL of 0.15 nmole dimer aptamers were incubated for 4 and 6 hours in RPMI-1640 with 10% FBS at 37°C with 5% CO_2_. At the end of each incubation time point, 1.2 µL of each aptamer were mixed with 8.8 µL of 1% TBE buffer and 2 µL of cyber green loading dye to make a dilution of 1:10. Samples along with low molecular weight ladder were loaded on a 4% Agarose gel and ran for 30 mins at 120 volts using Fisherbrand™ Electrophoresis power supplies (FB3000Q).

#### Cell Activation

Jurkat E6.1 cells were washed three times with 3 mL RPMI-1640 (+25mM HEPES, +L-Glutamine) medium prior to the activation study of TCR-CD3 in Jurkat E6.1 cell. Following cell wash, 2 × 10^5^ Jurkat E6.1 cells were incubated with 0.15 nmole of dimer aptamers or dimer random control along with 10 µL of 25 µg/ml unlabeled anti-human CD28 (BioXCells) in 150 µL CSB. The cells, dimer aptamers and the anti-human CD28 antibody were incubated for 2, 4, and 6 hours in a surface-modified polystyrene 96 well flat-bottom plate at 37°C with 5% CO_2_. This was followed by transferring the solution from the 96 well flat-bottom plate to a Ria Tubes 12 Χ 75 (Falcon ™ Round-Bottom polystyrene Tubes) to stain for CD69 receptor using 5 µL (100 µg/mL) of PreCP-Cy ™ 5.5 mouse anti-human CD69 (BD Biosciences). Staining was done for 45 mins on ice then washed one time with 2 mL RPMI-1640 medium and reconstituted in 250 µL RPMI-1640 medium. The expression of activation marker CD69 was monitored in FL4 Red (663-687 nM) at each time points using flow cytometry by counting 5000 events.

### Positive controls

#### Activation of cells with anti-CD3 and anti-CD28 antibody

Positive control experiments were performed by incubating cells with anti-human CD3ε and anti-human CD28 antibodies. The cells were washed three times with 3 mL RPMI-1640 (+25mM HEPES, +L-Glutamine) medium then 2 × 10^5^ Jurkat E6.1 cells were incubated for 2, 4 and 6 hours with 10 µL of 10 µg/mL unlabeled anti-human CD3ε and 10 µL of 25 µg/mL unlabeled anti-human CD28 antibodies at 37°C with 5% CO_2_ in a surface-modified polystyrene 96 well flat bottom plate. Cells were stained with 5 µL of 100 µg/mL CD69 mouse anti-human Cy5.5 antibody for 45 mins on ice then washed one time with 2 mL RPMI-1640 and reconstituted in 250 µL RPMI-1640 medium. The expression of activation marker CD69 was monitored in FL4 Red (663-687 nM) for each time points using flow cytometry by counting 5000 events.

#### Plate binding assay

A 96 well plate was first pre-coated with 50 µL (10 µg/mL) anti-CD3ε antibody for 2 hr at 37°C with 5% CO_2_. Following pre-coating, the 96 well plate was washed three times with sterile 1X PBS to remove the unbound antibodies. The cells were then prepared by washing two times with 2 mL RPMI-1640 medium. The 2 × 10^5^ Jurkat E6.1 cells were added to the plate along with 10 µL of 25 µg/mL anti-CD28 antibody at 37°C with 5% CO_2_ for 2, 4 and 6 hours. Following this, cells were stained with 5 µL of 100 µg/mL CD69 mouse anti-human Cy5.5 antibody for 45 mins on ice then washed one time with 2 mL RPMI-1640 and reconstituted in 250 µL RPMI-1640 medium. The expression of activation marker CD69 was monitored in FL4 for each time points using flow cytometry by counting 5000 events.

### Negative controls

#### Untreated cells in the absence of anti-CD3ε antibody or anti-CD28 antibody

Cells were prepared by washing three times with 3 mL RPMI-1640 medium. 2 × 10^5^ Jurkat E6.1 cells were incubated for 2, 4 and 6 hours in 150 µL cell suspension buffer in a surface-modified polystyrene 96 well flat bottom plate at 37°C with 5% CO_2_. After the incubation for 2, 4 and 6 hours, the cells were stained with 5 µL of 100 µg/mL CD69 mouse anti-human Cy5.5 antibody for 45 mins on ice. Following staining, cells were washed one time with 2 mL RPMI-1640 and reconstituted in 250 µL RPMI-1640 medium. The expression of activation marker CD69 was monitored in FL4 at each time points using flow cytometry by counting 5000 events.

#### Evaluating the decrease in activation of TCR-CD3 in Jurkat E6.1 cells in the presence of the dimers (6S, 8S) and the absence of anti-CD28 antibody

Cells were prepared by washing three times with 3 mL RPMI-1640 medium. 2 × 10^5^ Jurkat E6.1 cells were incubated with OSJ-D-6S or OSJ-D-8S dimer in 150 µL cell suspension buffer in the absence of anti-CD28 antibody at 37°C with 5% CO_2_ in a surface-modified polystyrene 96 well flat bottom plate. Cells were stained with 5 µL of 100 µg/mL CD69 mouse anti-human Cy5.5 antibody for 45 mins on ice then washed one time with 2 mL RPMI-1640 and reconstituted in 250 µL RPMI-1640 medium. The expression of activation marker CD69 was monitored in FL4 at each time points using flow cytometry by counting 5000 events.

#### Evaluating the lack of activation of TCR-CD3 in Jurkat E6.1 cells using the random dimer

Cells were prepared by washing three times with 3 mL RPMI-1640 medium. 2 × 10^5^ Jurkat E6.1 cells were incubated for 2, 4 and 6 hours with 0.15 nmole random dimer in 150 µL cell suspension buffer in the presence of 10 µL of 25 µg/mL unlabeled anti-CD28 at 37°C with 5% CO_2_ in a surface-modified polystyrene 96 well flat bottom plate. Following incubation, cells were stained with 5 µL of 100 µg/mL CD69 mouse anti-human Cy5.5 antibody for 45 mins on ice then washed one time with 2 mL RPMI-1640 and reconstituted in 250 µL RPMI-1640 medium. The expression of activation marker CD69 was monitored in FL4 at each time points using flow cytometry by counting 5000 events.

#### Evaluating the lack of activation of TCR-CD3 in Jurkat E6.1 cells using only anti-CD28 antibody

Cells were prepared by washing three times with 3 mL RPMI-1640 medium. 2 × 10^5^ Jurkat E6.1 cells were incubated for 2, 4 and 6 hours in the presence of 10 µL of 25 µg/mL unlabeled anti-CD28 antibody only at 37*°* C with 5% CO_2_ in a surface-modified polystyrene 96 well flat bottom plate. Following this, cells were stained with 5 µL of 100 µg/mL CD69 mouse anti-human Cy5.5 antibody for 45 mins on ice then washed one time with 2 mL RPMI-1640 and reconstituted in 250 µL RPMI-1640 medium. The expression of activation marker CD69 was monitored in FL4 at each time points using flow cytometry by counting 5000 events.

## Results

### Truncation and LNA −2’OMe RNA Incorporation

Recently, we introduced a new aptamer, ZUCH-1, (Figure 1A and Table S1) against the human CD3ε receptor in its native functional state using **Li**gand-**G**uided **S**election (LIGS), a variant of SELEX(16). LIGS identifies functional nucleic acid ligands against cell surface receptors in their native functional state from an evolved SELEX library. Thus, ZUCH-1 represents the first synthetic ligand against human TCR-CD3ε, a key receptor molecule in T-cell-mediated immune response. With dissociation constant in the nanomolar range, ZUCH-1 is suitable for designing aptamer-based T-cell activation agents. Thus, we sought to investigate the functionality of dimeric aptamers to activate TCR-CD3ε, while enhancing their affinity and stability. We strategically designed shorter variants of ZUCH-1 to further enhance the aptamer’s affinity without compromising its specificity. The optimized monomeric aptamer was further modified with nuclease-resistant RNA bases, followed by the design of a functional and stable dimeric aptamer against TCR-CD3ε.

**Figure 1:**
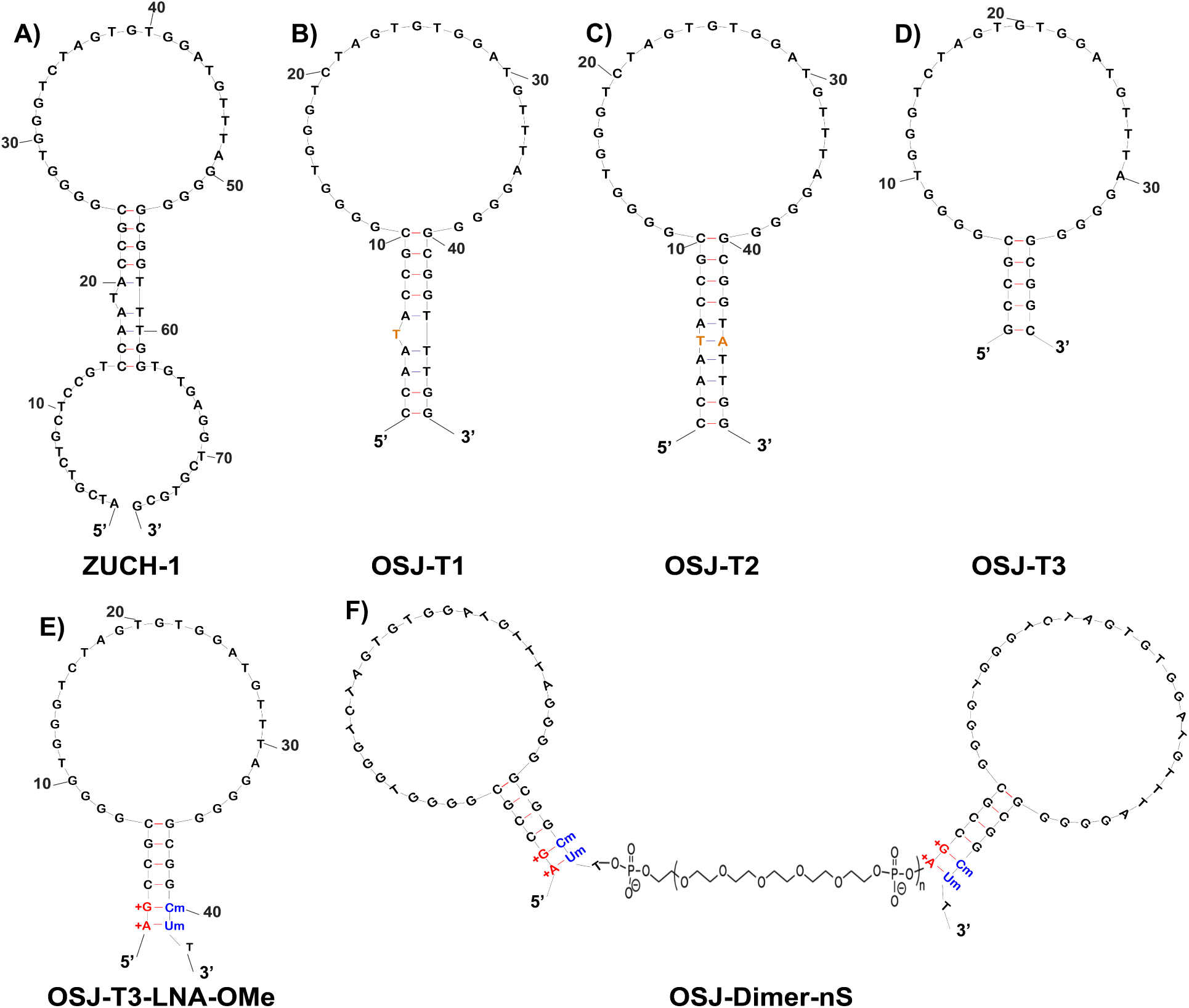
Truncation, modification, and dimerization of ZUCH-1 aptamer variants. A) Original ZUCH-1 aptamer. (B) Truncated OSJ-T1 variant, removing 14 bases from 5’ and 3’ terminals. (C) Stem-modified OSJ-T2 variant, adding adenosine at 45th position. (D) Truncated OSJ-T3 with the most optimized length with 39 nucleotides. (E) LNA- and 2’-OMe RNA-modified variant. (F) Dimerized variants of OSJ-T3-LNA-OMe with variable linkers to assess biological function, nS: number of Spacers.

The base length of the initially reported parent aptamer ZUCH-1 is 76 nt. However, as demonstrated by several studies, not all nucleotides in an aptamer play a significant role in aptamer folding, target recognition, or its function. Building on this assumption, we designed shorter variants of ZUCH-1. We first removed 14 nucleotides from ZUCH-1 at both the 5’ and 3’ terminal ends of the aptamer sequence, resulting in the OSJ-T1variant, as shown in Figure 1B. The calculated affinity of OSJ-T1 was similar to that of the full-length aptamer with a Kd of 2.370nM ±0.553 (Table 1 and Affinity curve in Figure 2A-(i)). The insignificant change in the affinity of OSJ-T1 suggests that bases removed from the terminal ends of the parent aptamer were nonessential and that their removal did not disrupt the functional fold critical for aptamer binding or recognition. However, OSJ-T1 consists of 48 nucleotides with one unpaired thymine base in the stem. Typically, unpaired bases in the stem region of a nucleic acid fold can destabilize the stem(22). To investigate whether stabilization of the stem improved affinity, we used OSJ-T1 as a template and modified the aptamer structure in a series of steps. First, we introduced a base modification by adding a complementary adenine to the 45^th^ base position in the stem of the OSJ-T1 sequence to remove the kink, resulting in variant OSJ-T2 (Figure 1C). However, with a calculated Kd of 2.761nM±0.4824 (Table 1), stabilization of the stem did not lead to higher affinity, suggesting that stem stabilization did not directly result in enhancing the overall functional structure of the aptamer, or the aptamer’s ability to recognize TCR-CD3 (Figure 2A-(ii)). Second, we further truncated the stem sequence of OSJ-T2 by removing a total of 10 bases from both terminal ends, which resulted in variant OSJ-T3, consisting of 39 nucleotides with an affinity of 2.185nM ± 0.2683 (Table1, Figure 1D and Figure 2A-(iii)).

**Table 1.** Affinities of truncated, modified and dimerized ZUCH-1 parent aptamer

| Name | Affinity Jurkat-ATCC (nM) |
| --- | --- |
| OSJ-T1 | 2.37±0.553 |
| OSJ-T2 | 2.761±0.4824 |
| OSJ-T3 | 2.185±0.268 |
| OSJ-T3-LNA-OMe | 1.722±0.037 |
| OSJ-Dimer-2S | 0.4855±0.09321 |
| OSJ-Dimer-4S | 0.2935±0.032.2 |
| OSJ-Dimer-6S | 0.4057±0.08135 |
| OSJ-Dimer-8S | 1.71±0.243 |

**Figure 2:**
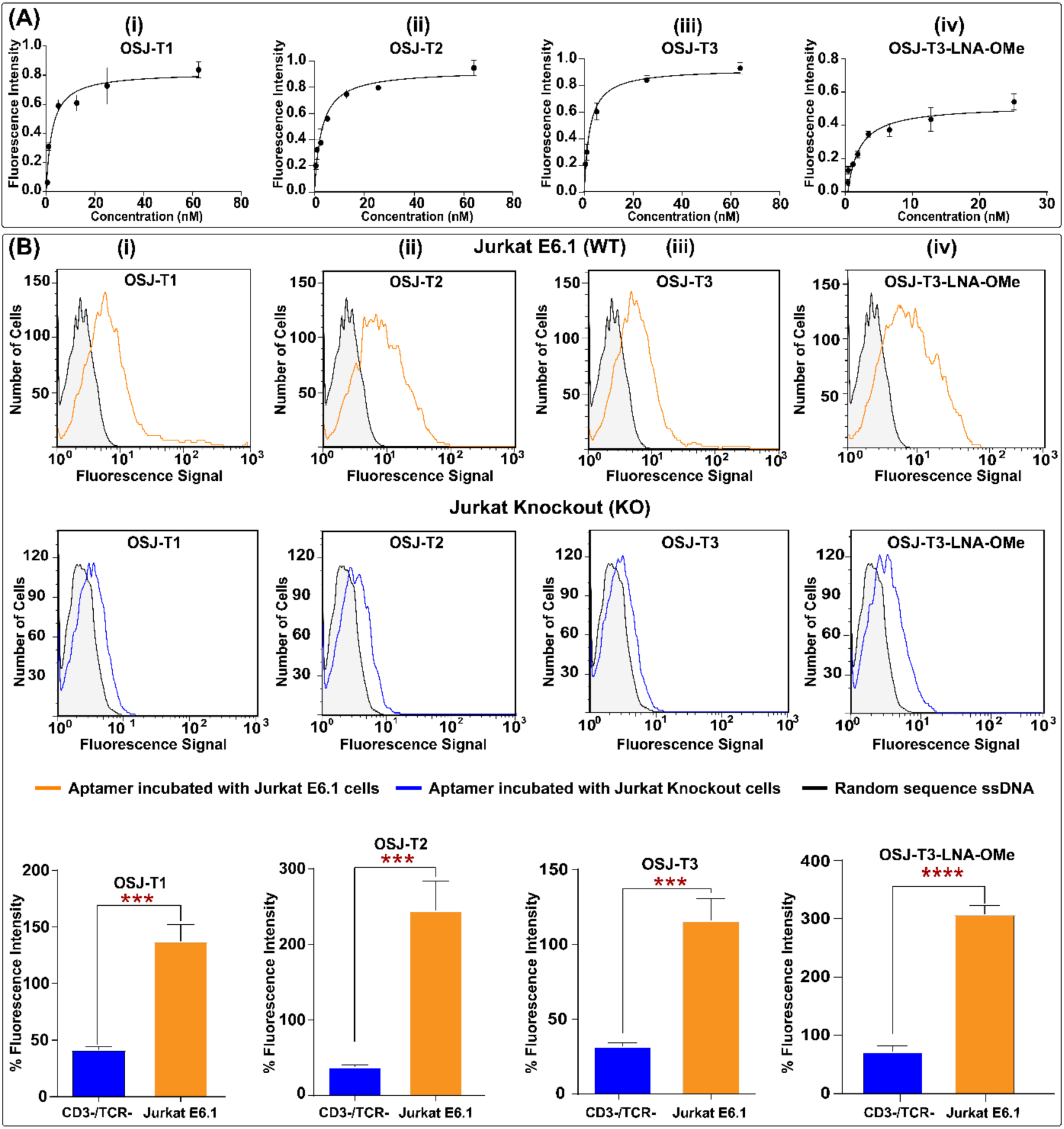
Analysis of affinity and specificity of truncated monomers and modified monomeric variants against TCR-CD3ε expressed on Jurkat E6.1 cells. (A) Affinity curves for truncated variants and modified monomer with unnatural RNA analogs were plotted, and the Kd of each was calculated using GraphPad Prism with a nonlinear fit, one site total and nonspecific binding. (B) The specificity of truncated variants and modified variant against TCR-CD3ε-positive Jurkat E6.1 cells (B, row 1) and TCR-CD3ε−negative engineered Jurkat E6.1 cells (B, row 2). Row 3 presents the overall conclusion of three independent specific binding experiments of the variant aptamers using one-way ANOVA with a t-test performed on GraphPad Prism ***:0.0004<p<0.0008, ****: p≤ 0.0001.

We next focused on increasing the robustness of the OSJ-T3 variant. We did this by improving nuclease resistance and structural stabilization by incorporating two different types of RNA analogs: Locked Nucleic Acids (LNA) and 2’OMe RNA bases (Figure 1E)(23). LNA bases are ribonucleotide (RNA) analogs containing a methylene link between the 2’-oxygen and 4’-carbon of the ribose ring(23). This constraint on the ribose moiety results in a locked C3’-endo hybridization that increases affinity towards its complementary base resulting in an enhancement of duplex stability(23,24). The 2’-O-methyl group (2’OMe RNA) is a naturally occurring modification found in RNA that enhances affinity for RNA-DNA targets owing to the preference of 2’-O-methyl-modified ribose sugars to adopt a C3’-endo conformation(25,26). These two unnatural bases have been shown to provide a significant improvement in duplex stability as a consequence of the high affinity of the two bases, leading to enhanced structural stabilization, stability against nucleases, and increased melting temperature(27-29). Previous reports have demonstrated that the high affinity of LNA and 2’OMe RNA analogs leads to superior duplex stability, both *in vitro* and *in vivo*(27). Nonetheless, aptamer post-SELEX modification with both LNA and 2’OMe RNA to enhance stability of a DNA aptamer has never been attempted. We predicted that modification with both LNA and 2’OMe RNA might lead to a highly stable aptamer in physiological conditions. Since RNA analogs lead to A-form RNA conformations, we systematically modified the variant OSJ-T3 with LNA and 2’OMe RNA bases to avoid introducing structural destabilization to OSJ-T3. To investigate the specific effects of LNA and 2’OMe RNA bases on the aptamer’s affinity, three variants of the OSJ-T3 aptamer were synthesized: 1) OSJ-T3-LNA, in which first and second bases from the 5’-end were modified with corresponding LNA bases (Figure S1, and 2(A)) OSJ-T3-OMe, in which the 40^th^ and 41^st^ bases from the 5’-end were modified with 2’OMe RNA bases (Figure S1, and 2(B)), and 3) OSJ-T3-LNA-OMe, with modifications to first and second bases from the 3’ terminal with LNA and 40^th^ and 41^st^ bases from the 5’-terminal base with 2’-OMe RNA bases, respectively (Figure 1E). As expected, aptamer OSJ-T3-LNA-OMe with dual modification provided heightened stabilization of the stem, leading to the formation of a functional secondary structure of the aptamer with a Kd value of 1.722nM± 0.037 ((Figure 2A-(iv), and Table 1).

### Specificity

The specificity of all truncated variants and modified monomers with RNA analogs were analyzed using the TCR-CD3-positive cell line Jurkat E6.1 and double knockout (DKO) negative cell line based on Jurkat.E6, which lacks TCR-CD3 complex (Figure 2B). The double knockout cell line, generated *via* a CRISPR/Cas9 system, lacked the TRAC gene that encodes the alpha constant chain of the TCR and the CD3ε gene that encodes the CD3ε polypeptide(30). The specificity of all truncated aptamers, OSJ-T1, OSJ-T2, and OSJ-T3 variants (Figure 2B (i-iii), panels (1-3)), was tested against positive cell line Jurkat E6.1 and negative cell line CD3ε double knockout (DKO) (Figure 2B (i, ii, iii). All aptamers were specific toward Jurkat E6.1 cells compared to the CD3ε double knockout (DKO) cells.

Interestingly, we observed that modification with either LNA or 2’OMe RNA at the terminal ends of the aptamer led to reduced specificity, despite careful optimization of the folding conditions. Thus, the resulting aptamers OSJ-T3-LNA and OSJ-T3-OMe did not show specific binding toward the TCR-CD3-expressing Jurkat cells (Figure S2). One way of explaining the observed loss of specificity of these two variants might be the undesirable hybridization of the modified bases to the natural bases on the hairpin, leading to the loss of the aptamer’s functional secondary structure. Undesirable fold formation owing to base modifications of the molecular beacon has been observed before(31,32). However, the dual-modified aptamer variant OSJ-T3-LNA-OMe, consisting of both LNA and 2’-OMe RNA base modifications at both terminals (Figure 1E), did maintain its specificity (Figure 2B (iv), fourth panel), suggesting that modifications at both ends did not interfere with the aptamer’s functional fold formation.

### Effect of different linker lengths on affinity and specificity

In addition to modification with unnatural bases, previous studies have also shown that the dimerization of aptamers can lead to increased binding affinity(18,20). We, therefore, dimerized the variant OSJ-T3-LNA-OMe with variable spacer lengths, consisting of phosphoramidite 18 with six glycol linkages (Figure 1E). We designed four dimeric arrangements with different spacer numbers, starting with two spacer modifiers consisting of 12 PEG units (2S), as the shortest, and eight spacer modifiers consisting of 48 PEG units (8S), as the longest. Two more dimers with six spacers (6S) and four spacers (4S) were also designed. We tested the binding affinity of all four dimers against Jurkat E.6.1 cells over a range of concentrations for each aptamer (Figure 3 A (i-iv)). The calculated Kd values (Table 1) were determined using GraphPad Prism software by plotting the specific median fluorescence intensity against the concentration of the aptamer variant. The binding affinities of the OSJ-D-2S, OSJ-D-4S, OSJ-D-6S, and OSJ-D-8S dimers showed Kd values of 0.4855nM±0.09321, 0.2935nM±0.032, 0.4057nM±0.08135, and 1.71±0.243, respectively (Table 1). The affinities exhibited by 2S, 4S, and 6S linkers averaged more than two-fold increase over the monomeric aptamer OSJ-T3-LNA-OMe, with a Kd=1.722nM±0.037, while the affinity of 8S was similar to that of the monomers.

**Figure 3:**
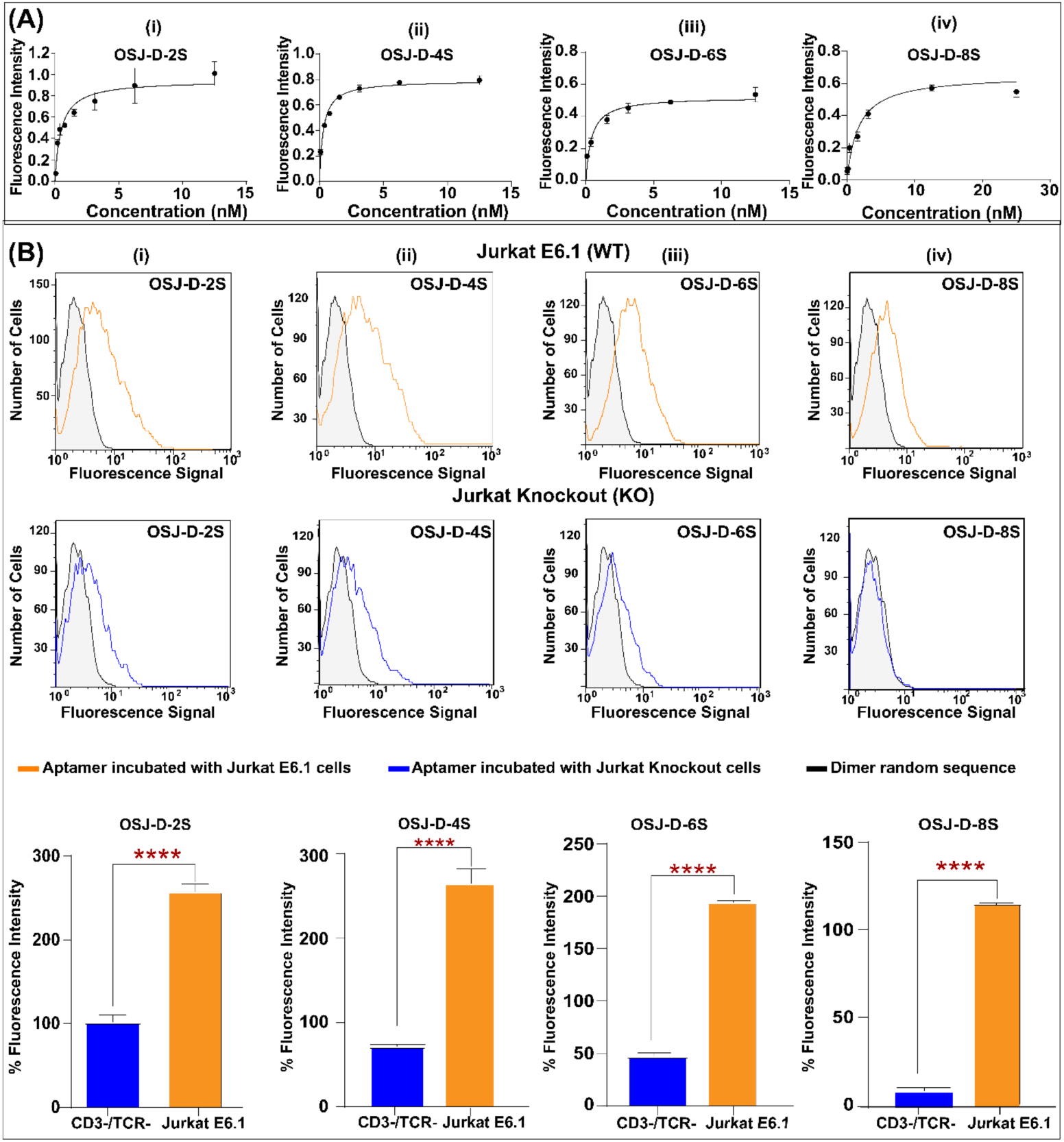
Analysis of affinity and specificity of dimeric variants against TCR-CD3ε expressed on Jurkat E6.1 cells. (A) Affinity curves for dimerized variants with 2 spacers consisting of 12 ethylene glycol units (i), four spacers with 24 ethylene glycol units (ii), six spacers with 36 ethylene glycol units (iii) and eight spacers with 48 ethylene glycol units (iv). Affinity curves were plotted, and the Kd of each was calculated using GraphPad Prism with a nonlinear fit, one site total and nonspecific binding. (B) Analysis of specificity of dimerized aptamer variants against TCR-CD3ε-positive wild-type (WT) Jurkat E6.1 cells (B, row 1) and CRISPR-Cas9 - engineered Jurkat E6.1 cells, knocking out (KO) TCR-CD3ε (B, row2). Row 3 shows the overall conclusion from three independent specificity analyses against KO and WT using one-way ANOVA and t-test test performed on GraphPad Prism ****: p≤ 0.0001.

We concluded that the observed increased affinities resulted from the spacing in the 2S, 4S, and 6S dimeric aptamer arrangements. Each TCR receptor possesses two epsilon domains, each capable of binding an aptamer domain(11). The dimeric aptamers with 2, 4, and 6 spacers may not be sufficient in length and/or flexibility for each aptamer in the dimeric scaffold to bind two different domains. However, the increase of local concentration could provide a tethered aptamer partner in close proximity that would have resulted in an increase in affinity. Besides, dimerization enhances the entropic penalty leading to highly favorable binding and, consequently, an increase in avidities in dimeric aptamer designs(33,34).

On the other hand, OSJ-D-8S showed affinity similar to that of the monomeric variant of aptamer OSJ-T3-LNA-OMe. Spacer length in the OSJ-D-8S dimer is approximately similar to the distance observed in the antibody arms of IgG antibodies. IgG antibodies are a broad family of antibodies, among which the OKT-3-specific anti-CD3ε monoclonal antibody was used in LIGS experiments to discover anti-TCR aptamers. Previous studies with microscopic imaging using OKT3 suggest that the distance between the two binding domains on the IgG antibody is approximately 150–180 Å(35-37). In the OSJ-D-8S dimer, the calculated distance between the two aptamers is approximately 168 Å, which falls between the optimal distances reported for an antibody(34,38,39). Similar dimensions between the OSJ-D-8S and OKT3 suggest that the two aptamer binding sites might interact with the two epsilon domains of the TCR-CD3 complex, leading to similar binding properties between OSJ-D-8S and an IgG-based OKT3 antibody.

Next, the nuclease resistance of all dimeric aptamers (OSJ-D-2S, OSJ-D-4S, OSJ-D-6S, and OSJ-D-8S) was investigated. Both LNA and 2’OMe-RNA modifications showed improved nuclease resistance. All dimeric aptamers were stable in 10% FBS (Figure 4) after 4 and 6 hours of incubation, suggesting that these modifications led to enhanced nuclease resistance of dimeric aptamers (Figure 4).

**Figure 4:**
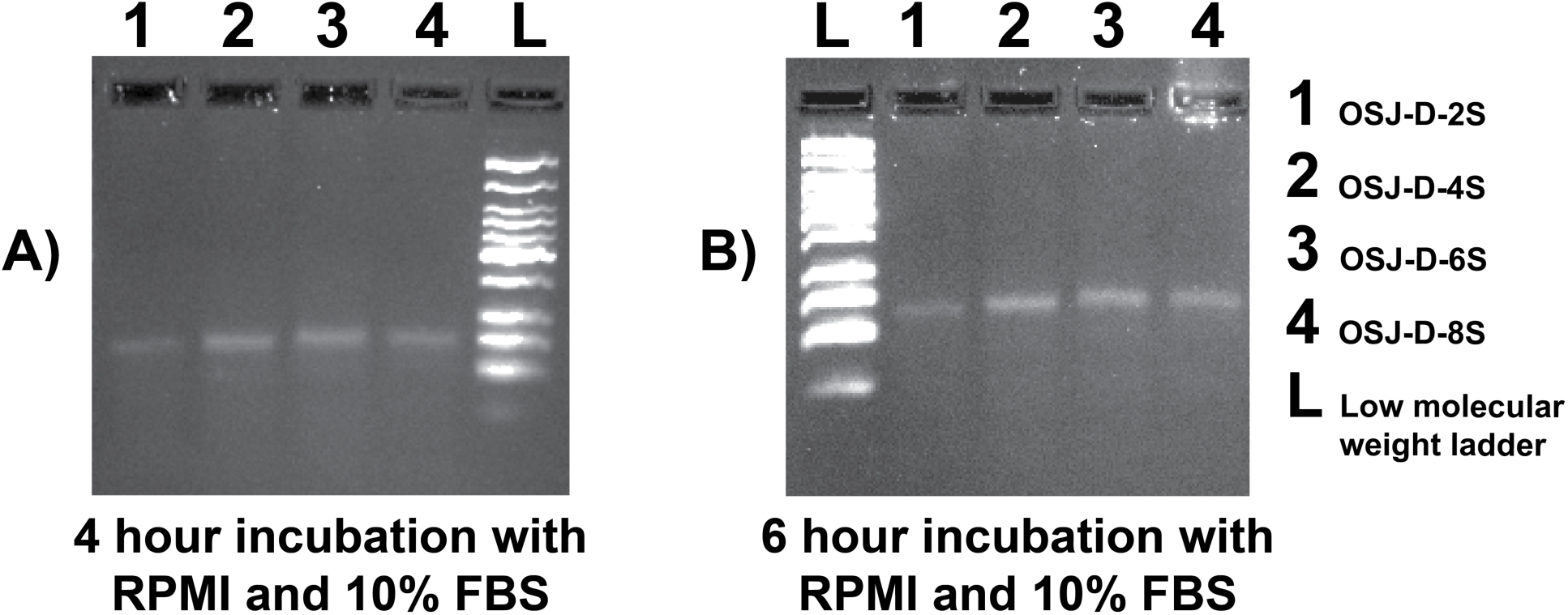
Analysis of nuclease resistance of LNA- and 2’OMe RNA-modified dimerized aptamers. (A) Analysis of 4 hours and (B) 6 hours incubation.

### Cell Activation from the 8S Dimer

Since the designed dimeric aptamer variants target TCR-CD3 complex, we next investigated whether a newly designed dimeric variant could activate TCR-CD3 on Jurkat.E6.1 cells. It is widely known that antibodies against CD3 can induce T-cell activation, and this concept has been used extensively in designing immunotherapeutic molecules(40). The antigenic activation of T cells leads to significant cellular changes essential to immune response. One of the earliest cell surface antigens expressed by T cells following activation is CD69, a membrane-bound, type II C-lectin receptor(41,42). CD69 expression is an early hallmark of lymphocyte activation based on its rapid appearance on the surface of the plasma membrane after stimulation(43). CD69 is not expressed on resting T-cells; therefore, we detected the activation of T-cells *via* the expression of CD69(43). We also monitored T-cell activation by the dimeric aptamer variants *via* the expression of CD69. Since the activation of Jurkat E 6.1 cells requires at least two signals to be fully activated, we used the anti-CD28 antibody as a costimulating agent(44,45). This activation was quantified using the CD69-labeled antibody (CD69 Mouse anti-Human Cy5.5). In the absence of anti-CD3 and anti-CD28, T-cells do not express CD69, confirming that both antibodies are required for activation hence expression of CD69 on the surface of T-cells (Figure S3). We then investigated whether the TCR-CD3 complex could be activated by all four dimeric aptamer variants, while using an anti-CD28 antibody as the secondary costimulator (Figure 5).

**Figure 5:**
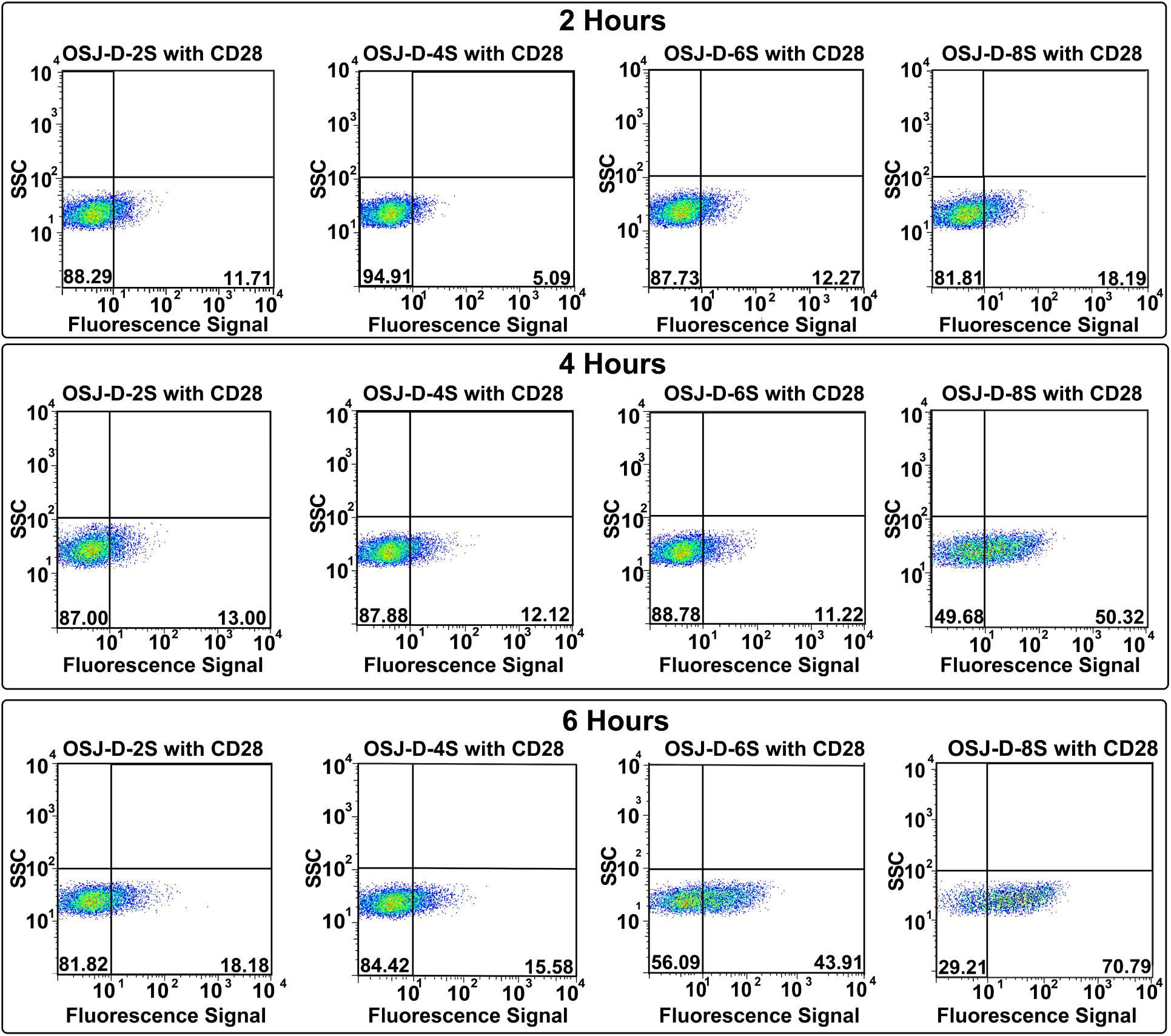
Investigation of the biological function of dimeric aptamer variants. The activation of TCR-CD3 in Jurkat E6.1 cells after incubation with 0.15 nmole dimer aptamers (D-2S, D-4S, D-6S, and D-8S, respectively) and 10 µL of 25 µg/mL unlabeled costimulatory anti-human CD28 antibody. The activation was studied for 2, 4 and 6 hours at 37°C with 5% CO_2_ and the appearance of early activation marker, CD69 receptor, were monitored using flow cytometry with 5 µL of 100 µg/mL anti-CD69 mAb-labeled with Cy5.5. The y-axis of the plots presents the side scatter of the cells (SSC). The x-axis is the fluorescence signal of anti-CD69 antibody. The contour plots correspond to each variant with variable spacers at three time points. The percentages beyond the cutoff line (bottom right quadrant) in the contour plots correspond to activated cells with positive expression of the CD69 receptor.

Jurkat E6.1 cells were incubated in cell suspension buffer for 2, 4, and 6 hours in a surface-modified polystyrene 96-well flat bottom plate with 0.15 nmoles of each dimeric aptamer and 25 µg/ml unlabeled anti-human CD28 at 37°C with 5% CO_2_. Following cell incubation of the dimeric aptamers, the cells were incubated with fluorescently labeled (Cy5.5) anti-CD69 antibody, followed by flow cytometric analysis. T-cell activation was calculated by comparing the number of cells expressing CD69 receptor compared to the number of cells undergoing no treatment (Figure S4). Cells expressing CD69 after 4 hours of incubation with the OSJ-D-8S dimeric aptamer suggested that OSJ-D-8S is an efficient activator of T-cells. However, the dimeric variants OSJ-D-2S, OSJ-D-4S, and OSJ-D-6S did not activate the cells and showed no expression of CD69. The OSJ-D-6S dimer did show a low level of activation after 6 hours. The positive control was included with the CD3ε antibody for 2, 4, and 6 hours in a surface-modified polystyrene 96-well flat bottom plate (Figure S4). Notably, the OSJ-D-8S aptamer showed activation of Jurkat E6.1 cells similar to that of the anti-human CD3ε antibody in 6 hours (Figure 5, third row)

To compare the activation with corresponding anti-CD3 antibody, a second positive control was performed using a 96-well plate pre-coated with 10 µg/ml CD3 for 2 hr at 37 °C and then washed twice, followed by adding the cells with CD28 antibody (Figure S4A and 4B). In addition, we performed multiple negative controls: (1) dimeric aptamer without anti-CD28 antibody, without costimulation, (2) randomized dimeric control with anti-CD28 antibody, in which aptamer-based stimulation was absent, while the costimulatory effect of anti-CD28 was present, and (3) anti-CD28 antibody with each experimental setup incubated for 2, 4, and 6 hours. All showed no T-cell activation suggesting that the activation observed for OSJ-D-8S was specific (Figure S5A-D)

## Discussion

Immunotherapeutic strategies, which involve directing T-cells towards diseased cells, such as cancer, have revolutionized drug development approaches(46,47). Currently, multiple immunotherapeutic approaches are being developed beyond cancer treatment. However, current approaches in developing immunotherapeutics are confined to either engineering T cells to express disease-targeting antigen in the form Chimeric Antigen Receptor T cells (CAR-T), or the use of engineered fragments of monoclonal antibodies (mAbs) in the form of bispecific antibodies or the use of mAbs against immune-checkpoint inhibitors(46-48). While all three strategies are successful in treating disease, particularly cancer, growing evidence suggests that immunotherapeutic approaches have their challenges. These challenges involve slow pharmacokinetics of mAbs, leading to immune-related-adverse-effects (iRAE), as well as the high cost of therapeutic development(49). This calls for low-cost, easily modifiable synthetic analogs with favorable pharmacokinetic properties, using alternative molecules to develop immunotherapeutics. In this regard, nucleic acid aptamers are promising alternative molecules. Aptamers are synthetic, easily modifiable, and show tunable pharmacokinetics(50,51). However, the challenge of using aptamers heavily relies on the discovery of aptamers capable of recognizing critical receptors on the immune cell surface. In particular, T-cell-receptor complex-CD3ε is one of the most sought after receptors to develop synthetic immunotherapeutics. We have successfully identified DNA aptamers against TCR-CD3 ε using a method we developed in our lab termed Ligand Guided Selection (LIGS). While we have discovered multiple aptamers against TCR-CD3 ε, aptamer ZUCH-1 showed the highest affinity. However, ZUCH-1 is 76 nucleotides in length, thus preventing the design of functional scaffolds.

Additionally, since natural nucleic acid bases are susceptible to nuclease degradation, it is necessary to modify aptamers with native bases with unnatural nucleic acid analogs. Thus, in this study, we successfully demonstrated that truncation followed by modification of unnatural nucleic acid analogs could further improve the parent aptamer without compromising its specificity, yet further enhancing its affinity. While improving the biostability of aptamers is an essential step in designing functional aptamers, an important goal of the current study is to explore the applicability of anti-TCR-CD3ε aptamers as T-cell activators. Out of all molecules tested, neither monomeric aptamers nor dimeric aptamers with shorter linkers activated TCR-CD3. However, the construct with eight spacers and dimensions similar to those of an antibody could activate TCR-CD3, suggesting that the design of functional dimeric aptamer scaffolds is predominantly governed by linker length, particularly against TCR-CD3ε.

Similarly, monovalent Fab fragment of anti-TCR antibody only poorly activated TCR-CD3, while the bivalent antibody could effectively activate TCR-CD3, suggesting that the dimeric aptamer analog against TCR-CD3ε might share activation characteristics similar to those of an antibody against the same receptor. A number of mechanisms of TCR-CD3 activation have been proposed. Still, the precise molecular interactions that explain how bivalent ligands, such as mAbs or dimeric aptamers, activate TCR-CD3 have not yet reached consensus, as indicated by the abundance of multiple controversial models(52,53). However, it has been shown that conformational change is required for optimal T-cell activation, suggesting that dimeric analog with eight spacers may induce a conformational switch to activate TCR-CD3.

In conclusion, we herein report the development of the first synthetic prototype of an anti-TCR-CD3 ligand, which was modified with LNA and 2’-methyl-RNA nucleic acid analogs. All constructs described here specifically recognize TCR-CD3 with high affinity. The observed differential activation properties suggest that dimensions of the bivalent design play a crucial role in activation of T-cell receptor complex. Future work will investigate the activation of TCR-CD3 using T-cells obtained from healthy volunteers and mechanistic investigation of TCR-CD3 activation by dimeric aptamers. To the best of our knowledge, no studies have, so far, reported on synthetic anti-TCR ligands capable of activating T-cells.

## Supplementary Data statement

Available online

## References

1. Weiner, L.M., Murray, J.C. and Shuptrine, C.W. (2012) Antibody-based immunotherapy of cancer. Cell, 148, 1081–1084.

2. Labrijn, A.F., Janmaat, M.L., Reichert, J.M. and Parren, P. (2019) Bispecific antibodies: a mechanistic review of the pipeline. Nature reviews. Drug discovery, 18, 585–608.

3. Sandigursky, S. and Mor, A. (2018) Immune-Related Adverse Events in Cancer Patients Treated With Immune Checkpoint Inhibitors. Curr Rheumatol Rep, 20, 65.

4. Belli, C., Zuin, M., Mazzarella, L., Trapani, D., D’Amico, P., Guerini-Rocco, E., Achutti Duso, B. and Curigliano, G. (2018) Liver toxicity in the era of immune checkpoint inhibitors: A practical approach. Crit Rev Oncol Hematol, 132, 125–129.

5. Hansel, T.T., Kropshofer, H., Singer, T., Mitchell, J.A. and George, A.J. (2010) The safety and side effects of monoclonal antibodies. Nature reviews. Drug discovery, 9, 325–338.

6. Dollins, C.M., Nair, S. and Sullenger, B.A. (2008) Aptamers in immunotherapy. Hum Gene Ther, 19, 443–450.

7. Pastor, F. (2016) Aptamers: A New Technological Platform in Cancer Immunotherapy. Pharmaceuticals (Basel), 9.

8. McNamara, J.O., Kolonias, D., Pastor, F., Mittler, R.S., Chen, L., Giangrande, P.H., Sullenger, B. and Gilboa, E. (2008) Multivalent 4-1BB binding aptamers costimulate CD8+ T cells and inhibit tumor growth in mice. J Clin Invest, 118, 376–386.

9. Dollins, C.M., Nair, S., Boczkowski, D., Lee, J., Layzer, J.M., Gilboa, E. and Sullenger, B.A. (2008) Assembling OX40 aptamers on a molecular scaffold to create a receptor-activating aptamer. Chem Biol, 15, 675–682.

10. Pastor, F., Soldevilla, M.M., Villanueva, H., Kolonias, D., Inoges, S., de Cerio, A.L., Kandzia, R., Klimyuk, V., Gleba, Y., Gilboa, E. et al. (2013) CD28 aptamers as powerful immune response modulators. Mol Ther Nucleic Acids, 2, e98.

11. Kuhns, M.S. and Badgandi, H.B. (2012) Piecing together the family portrait of TCR-CD3 complexes. Immunol Rev, 250, 120–143.

12. Tuerk, C. and Gold, L. (1990) Systematic evolution of ligands by exponential enrichment: RNA ligands to bacteriophage T4 DNA polymerase. Science, 249, 505–510.

13. Ellington, A.D. and Szostak, J.W. (1990) In vitro selection of RNA molecules that bind specific ligands. Nature, 346, 818–822.

14. Zumrut, H.E., Ara, M.N., Fraile, M., Maio, G. and Mallikaratchy, P. (2016) Ligand-Guided Selection of Target-Specific Aptamers: A Screening Technology for Identifying Specific Aptamers Against Cell-Surface Proteins. Nucleic Acid Ther, 26, 190–198.

15. Zumrut, H.E., Ara, M.N., Maio, G.E., Van, N.A., Batool, S. and Mallikaratchy, P.R. (2016) Ligand-guided selection of aptamers against T-cell Receptor-cluster of differentiation 3 (TCR-CD3) expressed on Jurkat.E6 cells. Anal Biochem, 512, 1–7.

16. Zumrut, H.E., Batool, S., Argyropoulos, K.V., Williams, N., Azad, R. and Mallikaratchy, P.R. (2019) Integrating Ligand-Receptor Interactions and In Vitro Evolution for Streamlined Discovery of Artificial Nucleic Acid Ligands. Mol Ther Nucleic Acids, 17, 150–163.

17. Zumrut, H., Yang, Z., Williams, N., Arizala, J., Batool, S., Benner, S.A. and Mallikaratchy, P. (2020) Ligand-Guided Selection with Artificially Expanded Genetic Information Systems against TCR-CD3epsilon. Biochemistry, 59, 552–562.

18. Mallikaratchy, P.R., Ruggiero, A., Gardner, J.R., Kuryavyi, V., Maguire, W.F., Heaney, M.L., McDevitt, M.R., Patel, D.J. and Scheinberg, D.A. (2011) A multivalent DNA aptamer specific for the B-cell receptor on human lymphoma and leukemia. Nucleic acids research, 39, 2458–2469.

19. Moccia, F., Riccardi, C., Musumeci, D., Leone, S., Oliva, R., Petraccone, L. and Montesarchio, D. (2019) Insights into the G-rich VEGF-binding aptamer V7t1: when two G-quadruplexes are better than one! Nucleic acids research, 47, 8318–8331.

20. Batool, S., Argyropoulos, K.V., Azad, R., Okeoma, P., Zumrut, H., Bhandari, S., Dekhang, R. and Mallikaratchy, P.R. (2019) Dimerization of an aptamer generated from Ligand-guided selection (LIGS) yields a high affinity scaffold against B-cells. Biochim Biophys Acta Gen Subj, 1863, 232–240.

21. Kimoto, M., Shermane Lim, Y.W. and Hirao, I. (2019) Molecular affinity rulers: systematic evaluation of DNA aptamers for their applicabilities in ELISA. Nucleic acids research, 47, 8362–8374.

22. Meroueh, M. and Chow, C.S. (1999) Thermodynamics of RNA hairpins containing single internal mismatches. Nucleic acids research, 27, 1118–1125.

23. Kaur, H., Babu, B.R. and Maiti, S. (2007) Perspectives on chemistry and therapeutic applications of Locked Nucleic Acid (LNA). Chem Rev, 107, 4672–4697.

24. Vester, B. and Wengel, J. (2004) LNA (locked nucleic acid): high-affinity targeting of complementary RNA and DNA. Biochemistry, 43, 13233–13241.

25. Iribarren, A.M., Sproat, B.S., Neuner, P., Sulston, I., Ryder, U. and Lamond, A.I. (1990) 2’-O-alkyl oligoribonucleotides as antisense probes. Proceedings of the National Academy of Sciences of the United States of America, 87, 7747–7751.

26. Beijer, B., Sulston, I., Sproat, B.S., Rider, P., Lamond, A.I. and Neuner, P. (1990) Synthesis and applications of oligoribonucleotides with selected 2’-O-methylation using the 2’-O-[1-(2-fluorophenyl)-4-methoxypiperidin-4-yl] protecting group. Nucleic acids research, 18, 5143–5151.

27. Mallikaratchy, P., Gardner, J., Nordstrom, L.U., Veomett, N.J., McDevitt, M.R., Heaney, M.L. and Scheinberg, D.A. (2013) A self-assembling short oligonucleotide duplex suitable for pretargeting. Nucleic Acid Ther, 23, 289–299.

28. Sun, B.W., Babu, B.R., Sorensen, M.D., Zakrzewska, K., Wengel, J. and Sun, J.S. (2004) Sequence and pH effects of LNA-containing triple helix-forming oligonucleotides: physical chemistry, biochemistry, and modeling studies. Biochemistry, 43, 4160–4169.

29. Yan, Y., Yan, J., Piao, X., Zhang, T. and Guan, Y. (2012) Effect of LNA- and OMeN-modified oligonucleotide probes on the stability and discrimination of mismatched base pairs of duplexes. J Biosci, 37, 233–241.

30. Cong, L., Ran, F.A., Cox, D., Lin, S., Barretto, R., Habib, N., Hsu, P.D., Wu, X., Jiang, W., Marraffini, L.A. et al. (2013) Multiplex genome engineering using CRISPR/Cas systems. Science, 339, 819–823.

31. Yang, C.J., Wang, L., Wu, Y., Kim, Y., Medley, C.D., Lin, H. and Tan, W. (2007) Synthesis and investigation of deoxyribonucleic acid/locked nucleic acid chimeric molecular beacons. Nucleic acids research, 35, 4030–4041.

32. Kim, Y., Yang, C.J. and Tan, W. (2007) Superior structure stability and selectivity of hairpin nucleic acid probes with an L-DNA stem. Nucleic acids research, 35, 7279–7287.

33. Meares, C.F. (2008) The chemistry of irreversible capture. Advanced drug delivery reviews, 60, 1383–1388.

34. Tian, L. and Heyduk, T. (2009) Bivalent ligands with long nanometer-scale flexible linkers. Biochemistry, 48, 264–275.

35. Saphire, E.O., Stanfield, R.L., Crispin, M.D., Parren, P.W., Rudd, P.M., Dwek, R.A., Burton, D.R. and Wilson, I.A. (2002) Contrasting IgG structures reveal extreme asymmetry and flexibility. J Mol Biol, 319, 9–18.

36. Saphire, E.O., Parren, P.W., Pantophlet, R., Zwick, M.B., Morris, G.M., Rudd, P.M., Dwek, R.A., Stanfield, R.L., Burton, D.R. and Wilson, I.A. (2001) Crystal structure of a neutralizing human IGG against HIV-1: a template for vaccine design. Science, 293, 1155–1159.

37. Zhang, X., Zhang, L., Tong, H., Peng, B., Rames, M.J., Zhang, S. and Ren, G. (2015) 3D Structural Fluctuation of IgG1 Antibody Revealed by Individual Particle Electron Tomography. Sci Rep, 5, 9803.

38. Xue, Y., O’Mara, M.L., Surawski, P.P., Trau, M. and Mark, A.E. (2011) Effect of poly(ethylene glycol) (PEG) spacers on the conformational properties of small peptides: a molecular dynamics study. Langmuir, 27, 296–303.

39. Clausen-Schaumann, H., Rief, M., Tolksdorf, C. and Gaub, H.E. (2000) Mechanical stability of single DNA molecules. Biophys J, 78, 1997–2007.

40. Liu, H., Rhodes, M., Wiest, D.L. and Vignali, D.A. (2000) On the dynamics of TCR:CD3 complex cell surface expression and downmodulation. Immunity, 13, 665–675.

41. Ziegler, S.F., Ramsdell, F. and Alderson, M.R. (1994) The activation antigen CD69. Stem Cells, 12, 456–465.

42. Cibrian, D. and Sanchez-Madrid, F. (2017) CD69: from activation marker to metabolic gatekeeper. Eur J Immunol, 47, 946–953.

43. Simms, P.E. and Ellis, T.M. (1996) Utility of flow cytometric detection of CD69 expression as a rapid method for determining poly- and oligoclonal lymphocyte activation. Clin Diagn Lab Immunol, 3, 301–304.

44. Molinero, L.L., Fuertes, M.B., Rabinovich, G.A., Fainboim, L. and Zwirner, N.W. (2002) Activation-induced expression of MICA on T lymphocytes involves engagement of CD3 and CD28. J Leukoc Biol, 71, 791–797.

45. Bashour, K.T., Gondarenko, A., Chen, H., Shen, K., Liu, X., Huse, M., Hone, J.C. and Kam, L.C. (2014) CD28 and CD3 have complementary roles in T-cell traction forces. Proceedings of the National Academy of Sciences of the United States of America, 111, 2241–2246.

46. Sadelain, M., Brentjens, R. and Riviere, I. (2013) The basic principles of chimeric antigen receptor design. Cancer Discov, 3, 388–398.

47. Inthagard, J., Edwards, J. and Roseweir, A.K. (2019) Immunotherapy: enhancing the efficacy of this promising therapeutic in multiple cancers. Clin Sci (Lond), 133, 181–193.

48. Brischwein, K., Parr, L., Pflanz, S., Volkland, J., Lumsden, J., Klinger, M., Locher, M., Hammond, S.A., Kiener, P., Kufer, P. et al. (2007) Strictly target cell-dependent activation of T cells by bispecific single-chain antibody constructs of the BiTE class. J Immunother, 30, 798–807.

49. Lemiale, V., Meert, A.P., Vincent, F., Darmon, M., Bauer, P.R., Van de Louw, A., Azoulay, E. and Groupe de Recherche en Reanimation Respiratoire du patient, d.O.-H. (2019) Severe toxicity from checkpoint protein inhibitors: What intensive care physicians need to know? Ann Intensive Care, 9, 25.

50. Batool, S., Bhandari, S., George, S., Okeoma, P., Van, N., Zumrut, H.E. and Mallikaratchy, P. (2017) Engineered Aptamers to Probe Molecular Interactions on the Cell Surface. Biomedicines, 5.

51. Keefe, A.D., Pai, S. and Ellington, A. (2010) Aptamers as therapeutics. Nature reviews. Drug discovery, 9, 537–550.

52. Iwashima, M. (2003) Kinetic perspectives of T cell antigen receptor signaling. A two-tier model for T cell full activation. Immunol Rev, 191, 196–210.

53. Minguet, S., Swamy, M., Alarcon, B., Luescher, I.F. and Schamel, W.W. (2007) Full activation of the T cell receptor requires both clustering and conformational changes at CD3. Immunity, 26, 43–54.

