## Supplemental file for "A homodimeric aptamer variant generated from ligand-guided selection activates T-cell receptor cluster of differentiation three complex"

**Table:** DNA sequences ZUCH1 and variants

| | |
|---|---|
| **ZUCH-1** | ATCGTCTGCTCCGTCCAATACCGCGGGGTGGGTCTAGTGTGGATGTTTAGGGGGCGGTTTG GTGTGAGGTCGTGCG |
| **OSJ-T1** | CCAATACCGCGGGGTGGGTCTAGTGTGGATGTTTAGGGGGCGGTTTGG |
| **OSJ-T2** | CCAATACCGCGGGGTGGGTCTAGTGTGGATGTTTAGGGGGCGGTATTGG |
| **OSJ-T3** | GCCGCGGGGTGGGTCTAGTGTGGATGTTTAGGGGGCGGC |
| **OSJ-T3-LNA** | +A+GCCGCGGGGTGGGTCTAGTGTGGATGTTTAGGGGGCGGC |
| **OSJ-T3-OMe** | AGCCGCGGGGTGGGTCTAGTGTGGATGTTTAGGGGGCGGCmUmT |
| **OSJ-T3-LNA-OMe** | +A+GCCGCGGGGTGGGTCTAGTGTGGATGTTTAGGGGGCGGCmUmT |
| **OSJ-D-2S** | +A+GCCGCGGGGTGGGTCTAGTGTGGATGTTTAGGGGGCGGCmUmT- $(C_{42}H_{61}N_2O_{10}P)_2$- +A+GCCGCGGGGTGGGTCTAGTGTGGATGTTTAGGGGGCGGCmUmT |
| **OSJ-D-4S** | +A+GCCGCGGGGTGGGTCTAGTGTGGATGTTTAGGGGGCGGCmUmT- $(C_{42}H_{61}N_2O_{10}P)_4$- +A+GCCGCGGGGTGGGTCTAGTGTGGATGTTTAGGGGGCGGCmUmT |
| **OSJ-D-6S** | +A+GCCGCGGGGTGGGTCTAGTGTGGATGTTTAGGGGGCGGCmUmT- $(C_{42}H_{61}N_2O_{10}P)_6$- +A+GCCGCGGGGTGGGTCTAGTGTGGATGTTTAGGGGGCGGCmUmT |
| **OSJ-D-8S** | +A+GCCGCGGGGTGGGTCTAGTGTGGATGTTTAGGGGGCGGCmUmT- $(C_{42}H_{61}N_2O_{10}P)_8$- +A+GCCGCGGGGTGGGTCTAGTGTGGATGTTTAGGGGGCGGCmUmT |

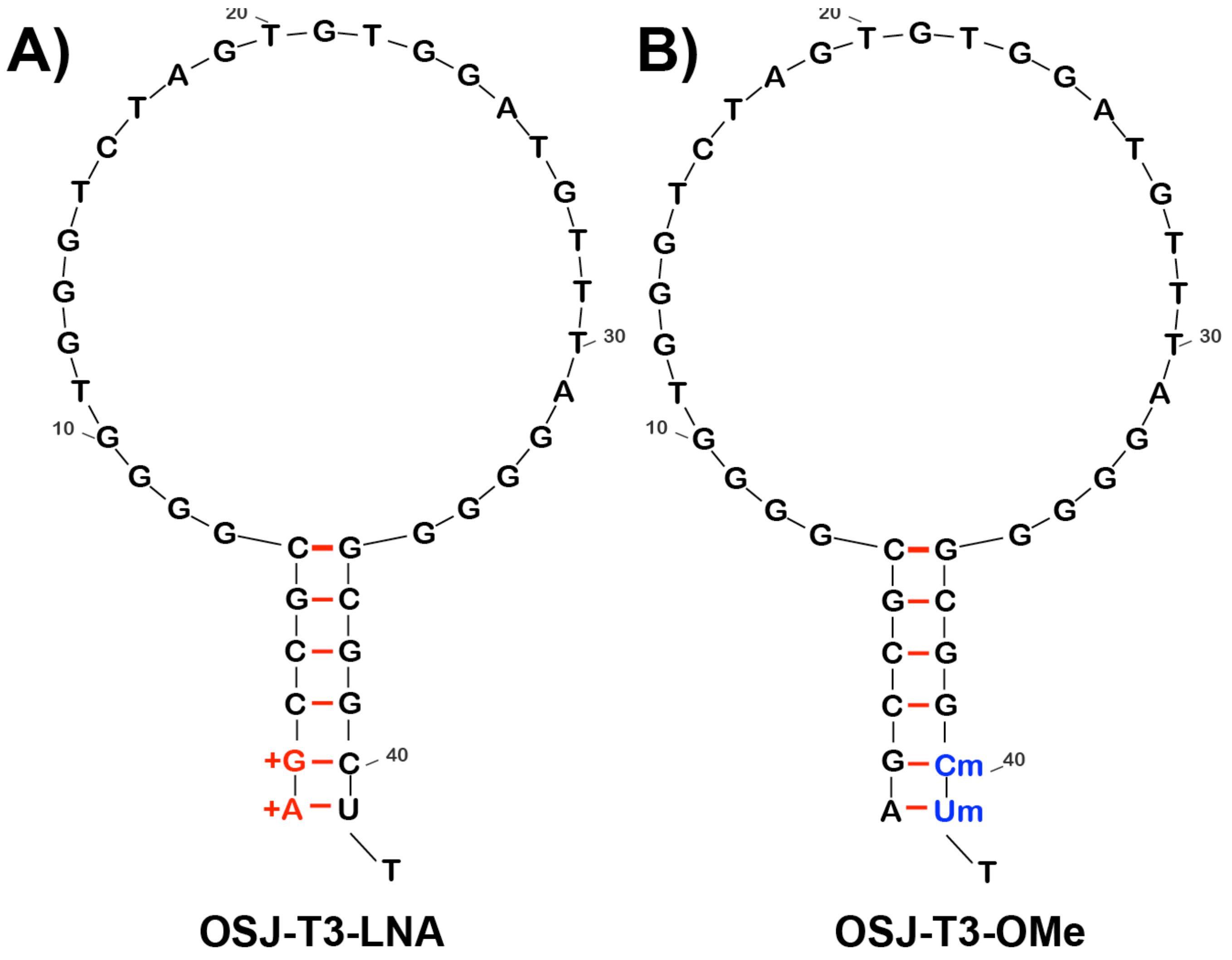


**Figure S1: The secondary structures of (A) OSJ-T3-LNA and (B) OSJ-T3-OMe modified with Locked Nucleic Acids (LNA) and 2'-O-methyl group (2'OMe RNA), respectively**. (A) bases modified within OSJ-T3-LNA sequence are indicated with a plus symbol, shown as "+", next to the base. (B) bases modified within OSJ-T3-OMe sequence are indicated with an "m" next to the base.

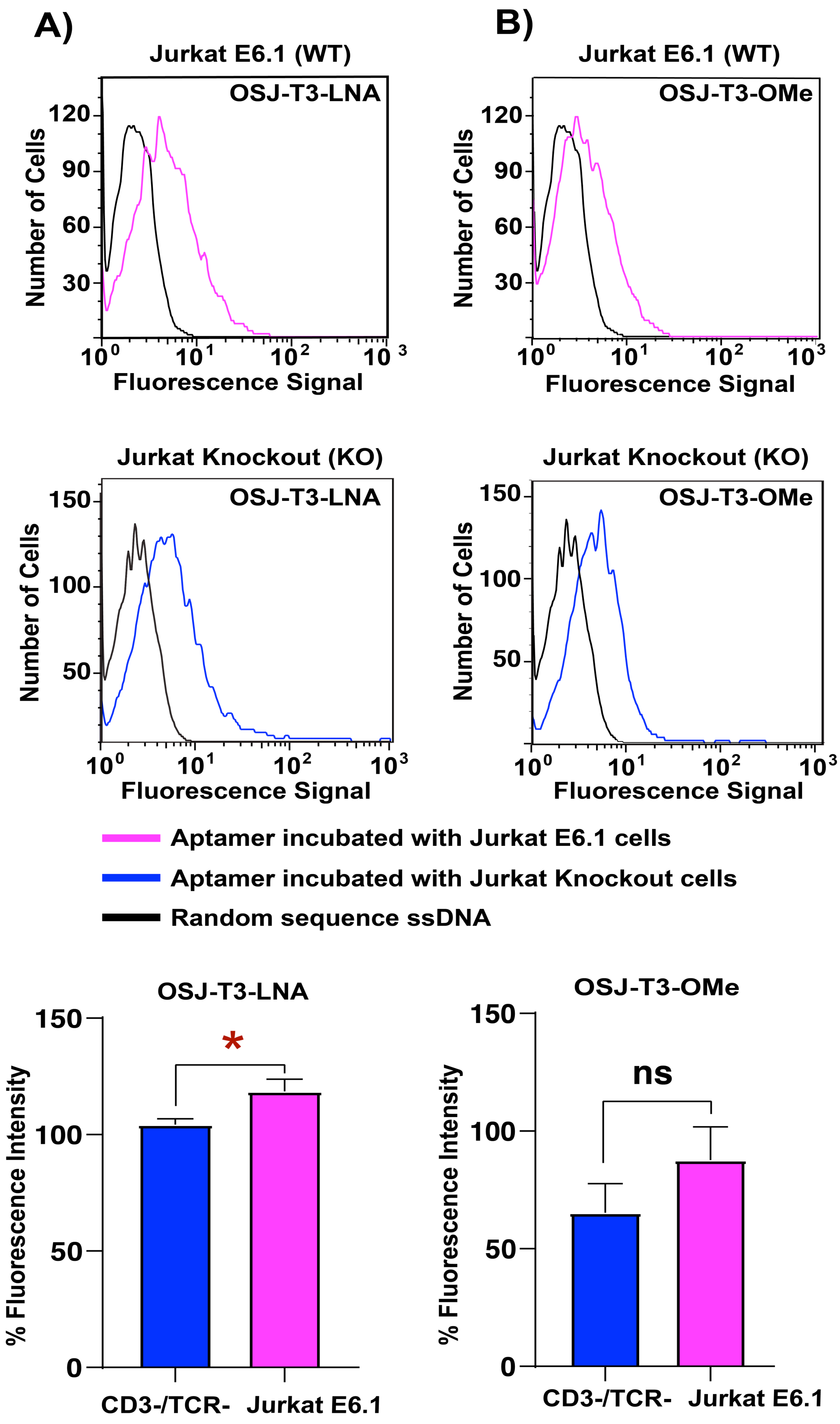


**Figure S2**: **Flow cytometry analysis of the specific binding of OSJ-T3-LNA and OSJ-T3-OMe towards Jurkat E6.1 WT (top row) and Jurkat CD3 knock-out cells (middle row)**. To investigate the specificity of OSJ-T3-LNA and OSJ-T3-OMe against Jurkat E6.1 WT cells, 0.0375 nmol of the 6-FAM-fluorophore labeled aptamers or random ssDNA molecules were denatured at 95°C for 10 mins and allowed to fold into its secondary structure at 4°C for 15 mins then 37°C for 30 mins. Aptamer or random molecules were then incubated with $1.5 \times 10^5$ Jurkat E6.1 WT or Jurkat CD3 knock-out cells in a total volume of 150 µL at 37°C. After incubation, the cells were washed two times with 2 mL of RPMI-1640 and reconstituted in 250 µL RPMI-1640 medium. Binding events were monitored in FL1 for the aptamer and random counting 5000 events using flow cytometry. (A) the specific binding of OSJ-T3-LNA towards Jurkat E6.1 WT (pink histogram) and Jurkat CD3 knockout cells (blue histogram). (B) the specific binding of OSJ-T3-OMe towards Jurkat E6.1 WT (pink histogram) and Jurkat CD3 knockout cells (blue histogram). Fluorescence intensity on the X axis represents FL1 shifts while the Y axis represents the number of cells. The specific binding for each aptamer was calculated by subtracting the random median fluorescence signal from the aptamer median fluorescence signal. The calculated specific binding from three independent experiments (for both WT and knockout cells) were used to calculate the specific binding percentages (*equation 1*). These percentages were then used to generate a bar graph using the one-way ANOVA with t-test performed on GraphPad Prism to obtain the statistical significance (third row). The bar graphs represent the overall three independent specific binding experiments of OSJ-T3-LNA and OSJ-T3-OMe. Both OSJ-T3-LNA and OSJ-T3-OMe did not show specific binding against Jurkat E6.1 WT cells. *: $p = 0.0140$

*Equation 1:*

$$\frac{\text{the median fluoresence signal of the aptamer} - \text{median fluoresence signal of the random}}{\text{median fluorescence signal of the random aptamer}} * 100$$

**Jurkat E6.1 cells untreated**

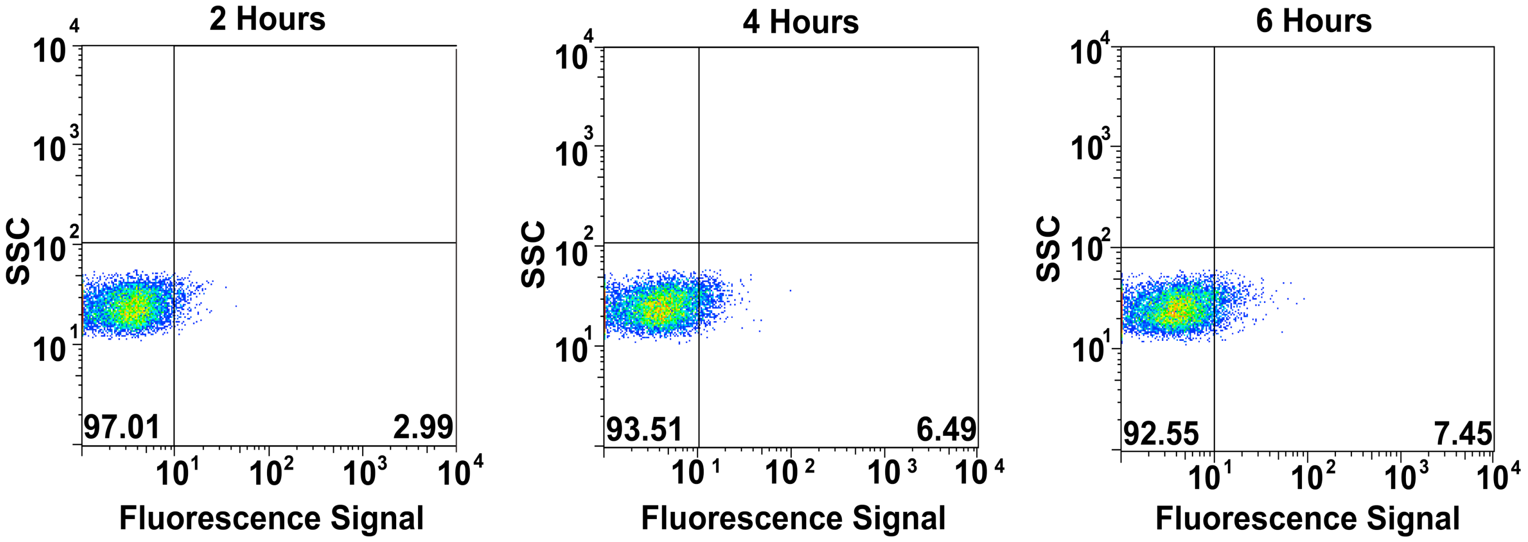


**Figure S3: Analysis of activation of TCR-CD3 in Jurkat E6.1 cell in the absence of unlabeled anti-CD3ε or anti-CD28 antibodies.** To study the activation of TCR-CD3 in Jurkat E6.1 cells, the cells were prepared by washing three times with 3 mL RPMI-1640 medium. $2 \times 10^5$ Jurkat E6.1 cells were incubated for 2, 4 and 6 hours in 150 µL cell suspension buffer in a surface-modified polystyrene 96 well flat bottom plate at 37°C with 5% $CO_2$. After the incubation for 2, 4 and 6 hours, the cells were stained with 5 µL of 100 µg/mL CD69 mouse anti-human Cy5.5 antibody for 45 mins on ice. Following staining, cells were washed one time with 2mL RPMI-1640 and reconstituted in 250 µL RPMI-1640 medium. The expression of activation marker CD69 was monitored in FL4 for each time points using flow cytometry by counting 5000 events. The fluorescence signal in FL4 vs. Side scatter (SSC) contour plots with quadrant gates showing four populations were generated using flow cytometry. The majority of cells incubated for 2, 4 and 6 hours (bottom left quadrant, 97.01%, 93.51% and 92.55%, respectively) did not express the activation marker CD69 even after 6 hours indicating no activation of TCR-CD3 in Jurkat E6.1 cell.

**Jurkat E6.1 cells treated**

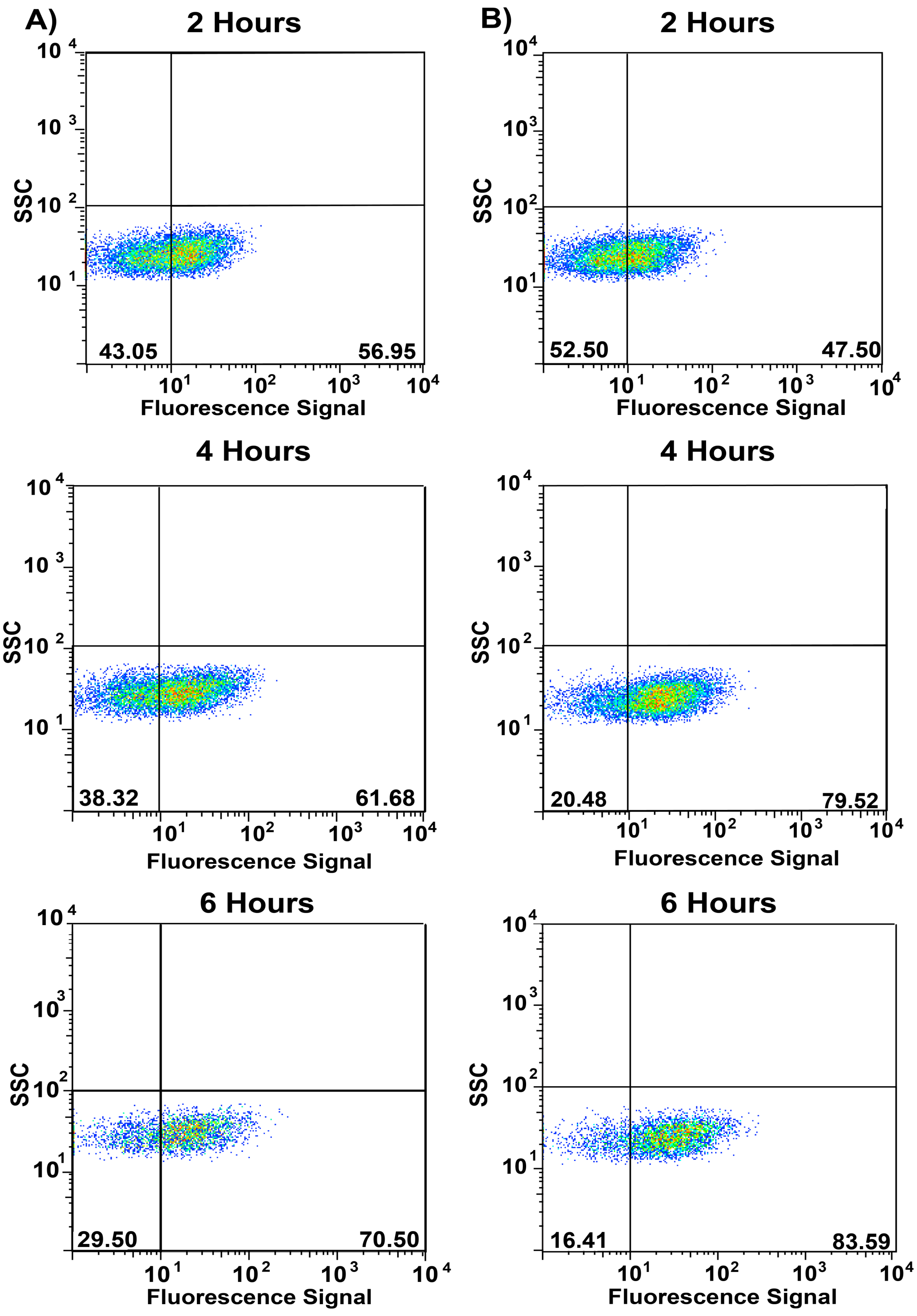


**Figure S4: Investigation of the activation of TCR-CD3 in Jurkat E6.1 cell in the presence of unlabeled anti-CD3ε and anti-CD28 antibodies.** Positive control experiments were performed by incubating cells with anti-human CD3ε and anti-human CD28 antibodies. (A) Cells were prepared by washing three times with 3 mL

RPMI-1640 medium. 2 × $10^5$ Jurkat E6.1 cells were incubated for 2, 4 and 6 hours with 10 µL of 10 µg/mL unlabeled anti-human CD3ε and 10 µL of 25 µg/mL unlabeled anti-human CD28 antibodies at 37 ° C with 5% $CO_2$ in a surface-modified polystyrene 96 well flat bottom plate. Cells were stained with 5 µL of 100 µg/mL CD69 mouse anti-human Cy5.5 antibody for 45 mins on ice and washed one time with 2 mL RPMI-1640 and reconstituted in 250 µL RPMI-1640 medium. The expression of activation marker CD69 was monitored in FL4 for all time points using flow cytometry by counting 5000 events. The fluorescence signal in FL4 vs. Side scatter (SSC) contour plots showed a gradual increase in the CD69 positive population for 2, 4 and 6 hours (bottom right quadrant, 56.95%, 61.68% and 70.50%, respectively) and a gradual decrease in the CD69 negative population for 2, 4 and 6 hours (bottom left quadrant, 43.05%, 38.32% and 29.50%, respectively). (B) a 96 well plate was first pre-coated with 50 µL (10 µg/mL) unlabeled anti-CD3ε antibody for 2 hours at 37°C with 5% $CO_2$ then washed three times with sterile 1X PBS. Following pre-coating, cells were washed two times with 2 mL RPMI-1640 and the 2 × $10^5$ Jurkat E6.1 cells were added to the plate for each time point along with 10 µL of 25 µg/mL anti-CD28 antibody at 37° C with 5% $CO_2$. Cells were stained with 5 µL of 100 µg/mL CD69 mouse anti-human Cy5.5 antibody for 45 mins on ice then washed one time with 2 mL RPMI-1640 and reconstituted in 250 µL RPMI-1640. The expression of activation marker CD69 was monitored in FL4 for all time points using flow cytometry by counting 5000 events. The fluorescence signal in FL4 vs. Side scatter (SSC) contour plots showed a gradual increase in the CD69 positive population for 2, 4 and 6 hours (bottom right quadrant, 47.50%, 79.52% and 83.59%, respectively) and a gradual decrease in the CD69 negative population for 2, 4 and 6 hours (bottom left quadrant, 52.50%, 20.48% and 16.41%, respectively). The overall expression of activation marker CD69 increased with longer incubation time.

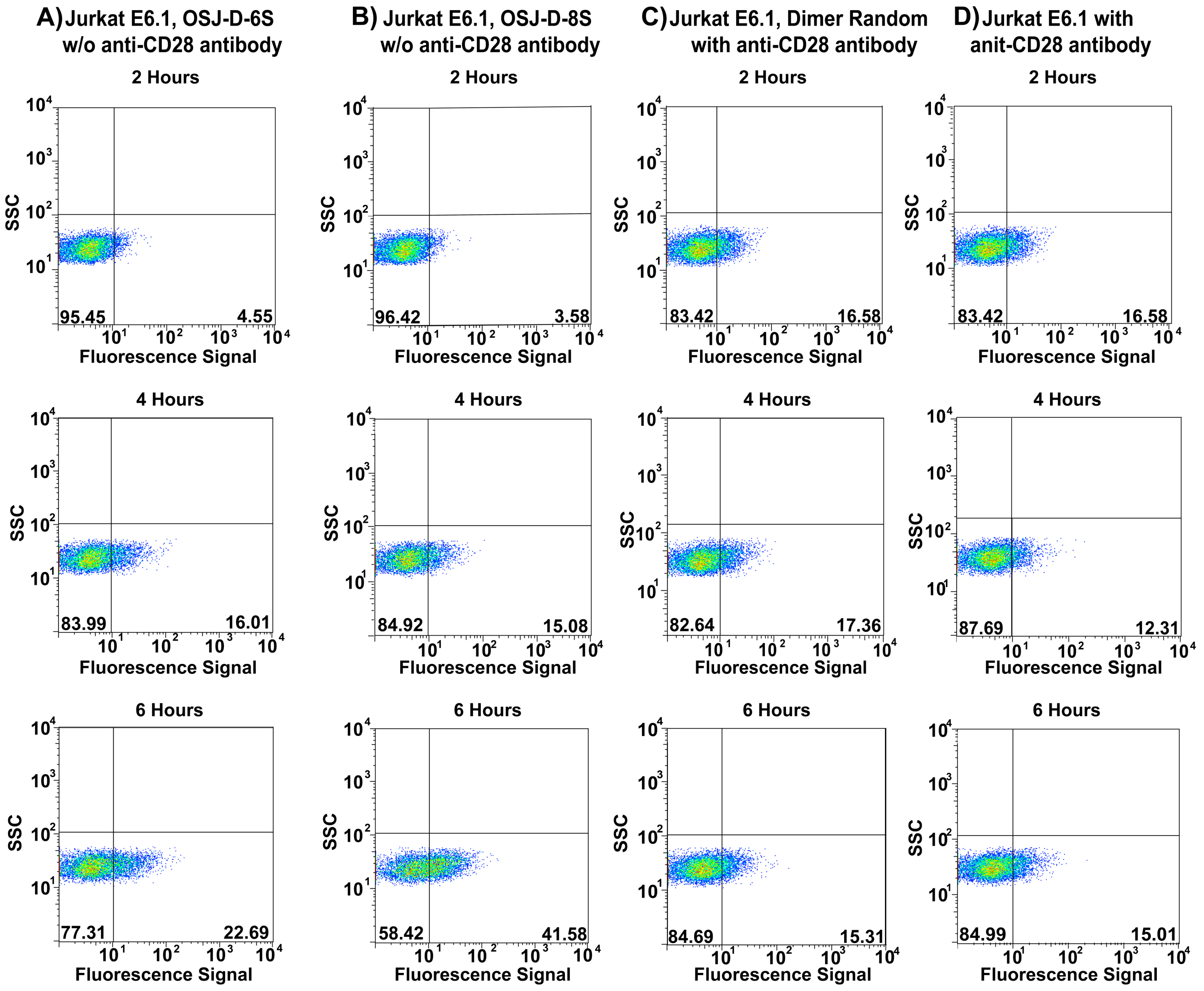


**Figure S5: Investigation of the activation of TCR-CD3 in Jurkat E6.1 cells with OSJ-D-6S and OSJ-D-8S dimeric aptamers in the presence or absence of anti-CD28 antibody.** Negative control experiments were performed by washing Jurkat E6.1 cells three times with RPMI-1640 medium then incubating the cells with OSJ-D-6S or OSJ-D-8S dimer in the absence of unlabeled anti-CD28 antibody (A-B) and random dimer in the presence of 10 μL of 25 μg/mL unlabeled anti-human CD28 antibody (C) or cells only in the presence of 10 μL of 25 μg/mL unlabeled anti-human CD28 antibody (D). (A) $2 \times 10^5$ Jurkat E6.1 cells were incubated for 2, 4 and 6 hours with 0.15 nmole OSJ-D-6S dimer in 150 μL cell suspension buffer in the absence of unlabeled anti-CD28 antibody at 37 °C with 5% $CO_2$ in a surface-modified polystyrene 96 well flat bottom plate. Cells were stained with 5 μL CD69 mouse anti-human Cy5.5 antibody for 45 mins on ice then washed one time with 2 mL

RPMI-1640 and reconstituted in 250 μL RPMI-1640 medium. (B) 2 × $10^5$ Jurkat E6.1 cells were incubated for 2, 4 and 6 hours with 0.15 nmole OSJ-D-8S dimer in 150 μL cell suspension buffer in the absence of unlabeled anti-human CD28 antibody at 37 ° C with 5% $CO_2$ in a surface-modified polystyrene 96 well flat bottom plate. Cells were stained with 5 μL of 100 μg/mL CD69 mouse anti-human Cy5.5 antibody for 45 mins on ice then washed one time with 2 mL RPMI-1640 and reconstituted in 250 μL RPMI-1640 medium. (C) 2 × $10^5$ Jurkat E6.1 cells were incubated for 2, 4 and 6 hours with 0.15 nmole random dimer in 150 μL cell suspension buffer in the presence of 10 μL unlabeled anti-CD28 antibody (25 μg/mL) at 37 ° C with 5% $CO_2$ in a surface-modified polystyrene 96 well flat bottom plate. Cells were stained with 5 μL of 100 μg/mL CD69 mouse anti-human Cy5.5 antibody for 45 mins on ice then washed one time with 2 mL RPMI-1640 and reconstituted in 250 μL RPMI-1640 medium. (D) 2 × $10^5$ Jurkat E6.1 cells were incubated for 2, 4 and 6 hours in the presence of 10 μL unlabeled anti-CD28 antibody (25 μg/mL) only at 37 ° C with 5% $CO_2$ in a surface-modified polystyrene 96 well flat bottom plate. Cells were stained with 5 μL of 100 μg/mL CD69 mouse anti-human Cy5.5 antibody for 45 mins on ice then washed one time with 2 mL RPMI-1640 and reconstituted in 250 μL RPMI-1640 medium. The expression of activation marker CD69 was monitored in FL4 for all time points using flow cytometry by counting 5000 events.  The fluorescence signal in FL4 vs. Side scatter (SSC) contour plots showed the majority of cells incubated for 2, 4 and 6 hours for all negative control experiments did not express the activation marker CD69, except for cells incubated with OSJ-D-8S for 6 hours (B, forth row) which showed a slight increase in CD69 positive population (bottom right quadrant, 41.58%). The OSJ-D-8S dimer did not activate the cells at the same level of activation as with anti-CD28 antibody. This data indicates that the activation of TCR-CD3 in Jurkat E6.1 cell requires co-stimulation of both the CD3ε receptor and the CD28 receptor.
